## Supplementary material for "The white-footed deermouse, an infection-tolerant reservoir for several zoonotic agents, tempers interferon responses to endotoxin in comparison to the mouse and rat": Figure S6

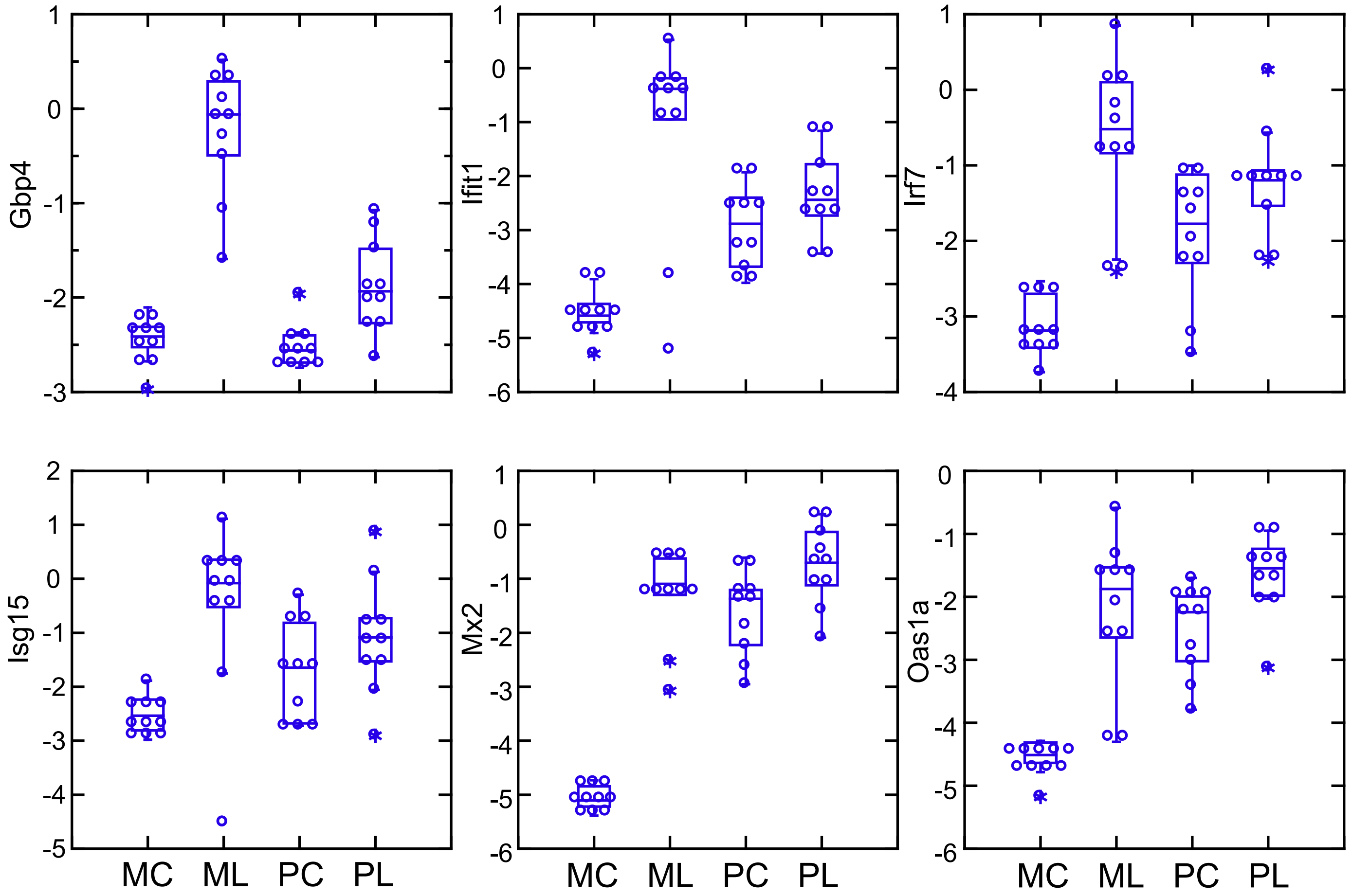


Figure S6. Box plots of log-transformed normalized transcripts for six interferon-stimulated genes of *Peromyscus leucopus* (P) or *Mus musculus* (M) that have been treated with LPS (L) or were saline-alone controls (C). There were 10 animals in each group and equally divided between females and males. Blood was obtained 4 hours after injection of LPS or saline alone, as described in Methods. Unique reads were normalized for reads of Ptprc for the same species for a given sample and the natural logarithm of the ratio was calculated.
