## Supplementary material for "The white-footed deermouse, an infection-tolerant reservoir for several zoonotic agents, tempers interferon responses to endotoxin in comparison to the mouse and rat": Figure S7

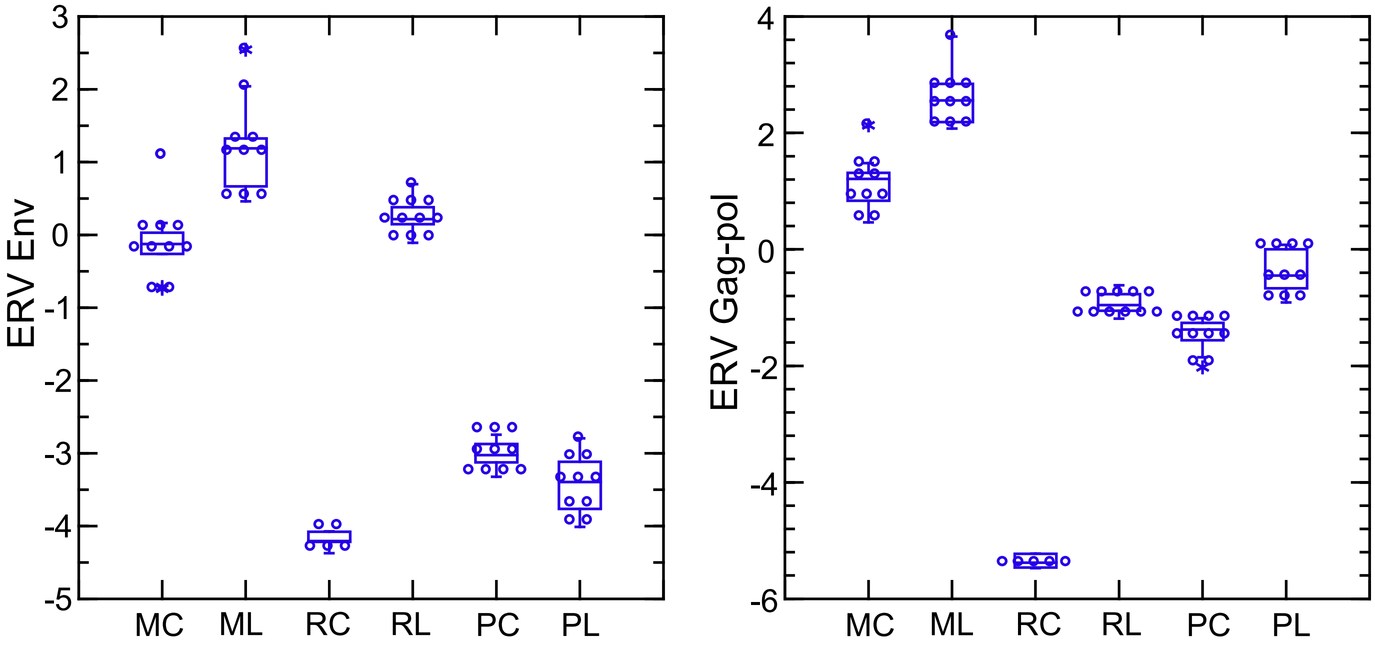


Figure S7. Box plots of log-transformed normalized transcripts in blood of envelope protein (Env) and Gag-pol protein chomosomal sequences of endogenous retroviruses (ERV) of *Peromyscus leucopus* (P), *Mus musculus* (M), or *Rattus norvegicus* (R) that have been treat-ed with LPS (L) or were saline-alone controls (C). Because of length differences for these coding sequences between 3 species, the unit used for cross-species analysis was reads per kilobase before normalization for Ptprc transcription.
