## Supplementary material for "The white-footed deermouse, an infection-tolerant reservoir for several zoonotic agents, tempers interferon responses to endotoxin in comparison to the mouse and rat": Figure S4

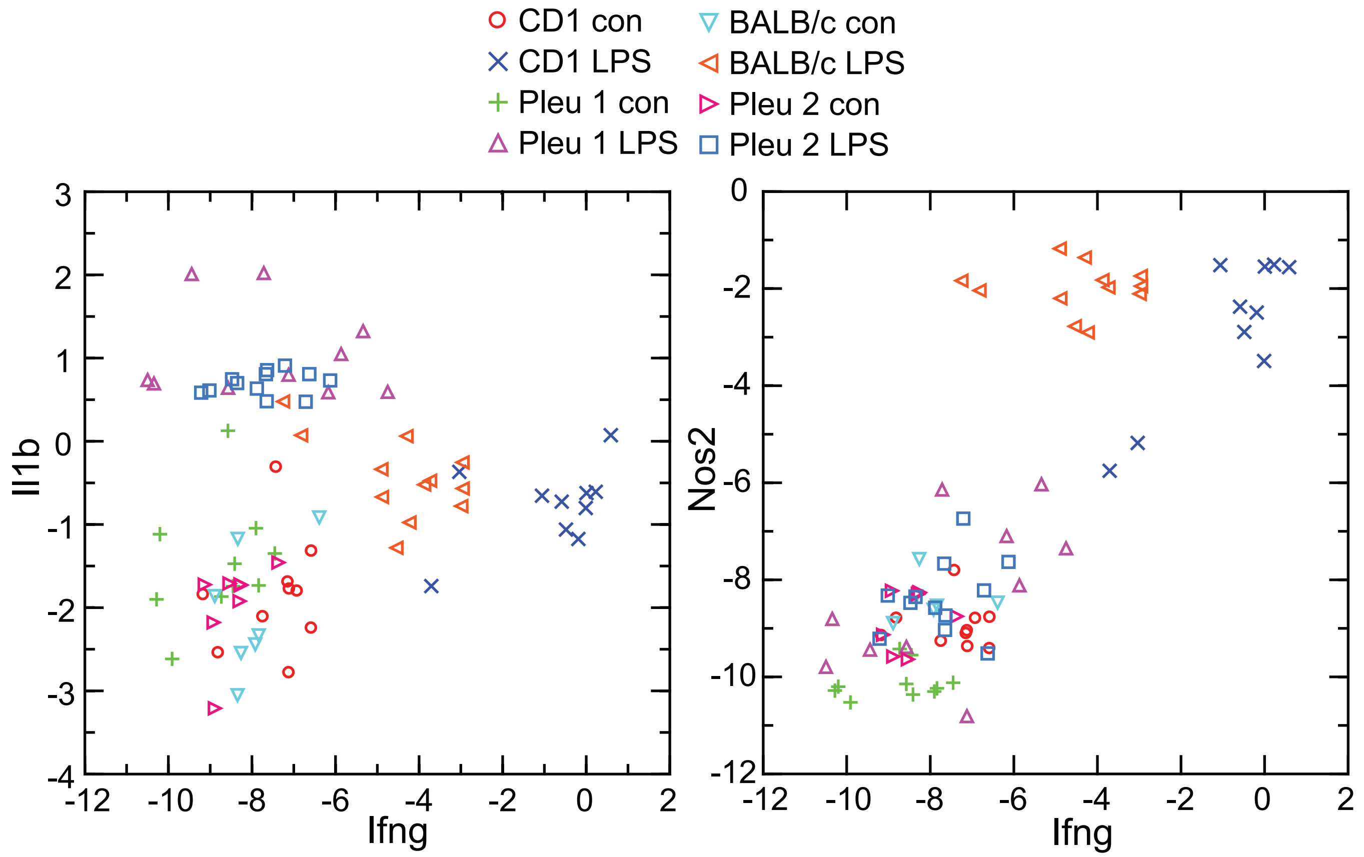


Figure S4. Scatter plots of log-transformed normalized transcripts of interleukin-1 beta (Il1b; left panel) or nitric oxide synthase 2 (Nos2; right panel) on interferon-gamma (Ifng) of blood of *P. leucopus* (Pleu) or *M. musculus* (outbred CD-1 and inbred BALB/c) with (LPS) or without (con) treatment with lipopolysaccharide 4 h previously. The data are from the present study (Pleu 2 and CD-1) and a further analysis of data (Pleu 1 and BALB/c) from the study of Balderrama-Gutierrez et al. (reference 3).
