## Supplementary material for "The white-footed deermouse, an infection-tolerant reservoir for several zoonotic agents, tempers interferon responses to endotoxin in comparison to the mouse and rat": Figure S3

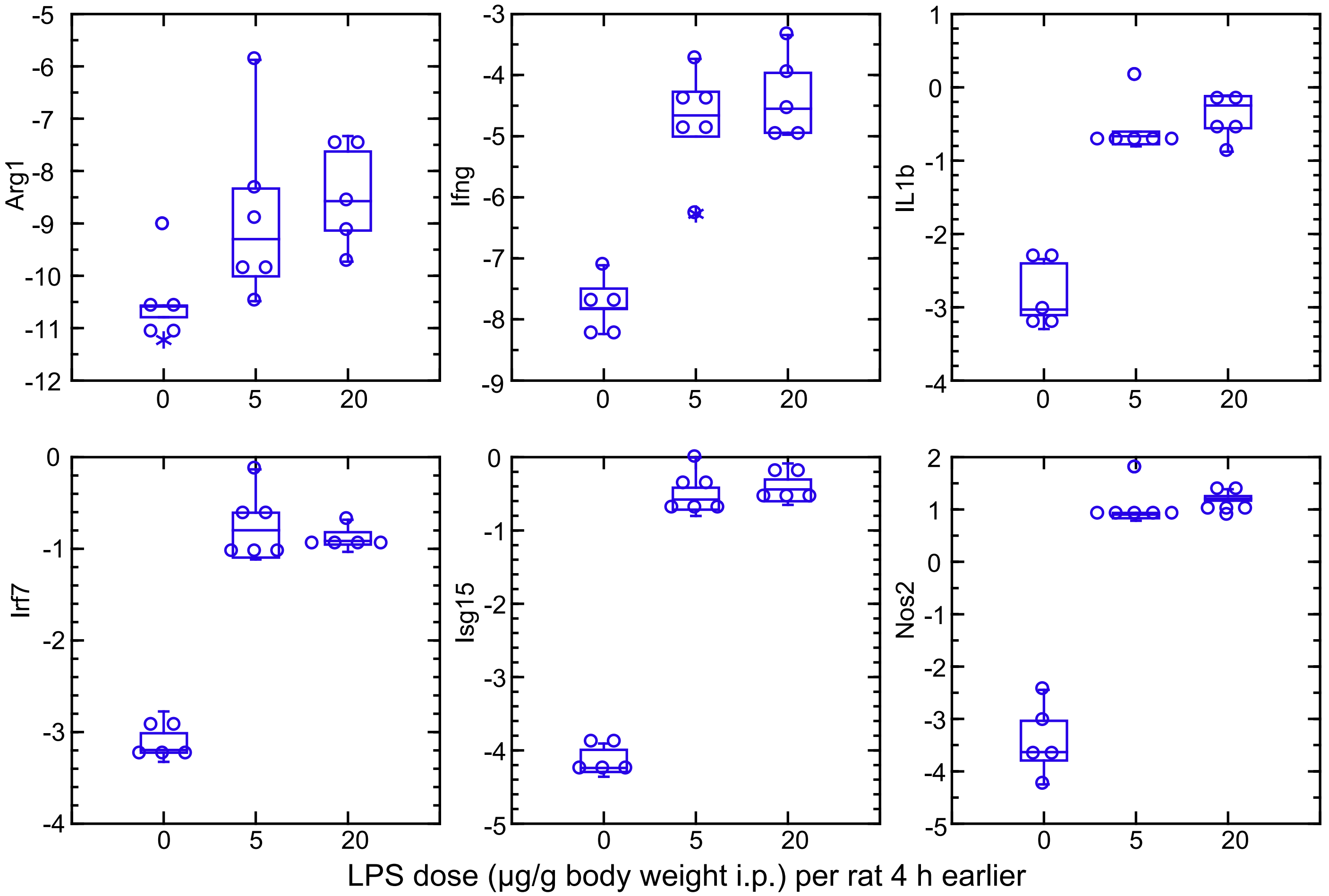


Figure S3. Box plots of log-transformed normalized transcripts of blood for six genes of adult female Rattus norvegicus (rat) injected intraperitoneally (i.p.) with saline alone (n = 5) or either of two doses LPS: 5 µg/g body weight (n = 5), or 20 µg per g body weight (n = 6). Blood was obtained 4 h after injection.
