## Supplementary material for "The white-footed deermouse, an infection-tolerant reservoir for several zoonotic agents, tempers interferon responses to endotoxin in comparison to the mouse and rat": Figure S2

Figure S2. Box plots of log-transformed normalized transcripts in whole blood for 54 genes of *Peromyscus leucopus* (P) or *Mus musculus* (M) that have been treated with LPS (L) or were saline-alone controls (C). There were 10 animals in each group and equally divided between females and males. Blood was obtained 4 hours after injection of LPS or saline alone, as described in Methods. Unique reads were normalized for reads of Ptprc for the same species for a given sample, and natural logarithm (*ln*) of the ratio calculated. These genes were drawn from the list of Table 2 as representative of different outcomes within the following general categories and symbols in the figure and, in most cases, not given more in-depth attention elsewhere in the study.

- ● metabolism
- ● kinases and related
- ● white cell markers
- ● proteins in plasma
- ● transcription factors
- ● neutrophil associated
- ● cytokines/chemokines

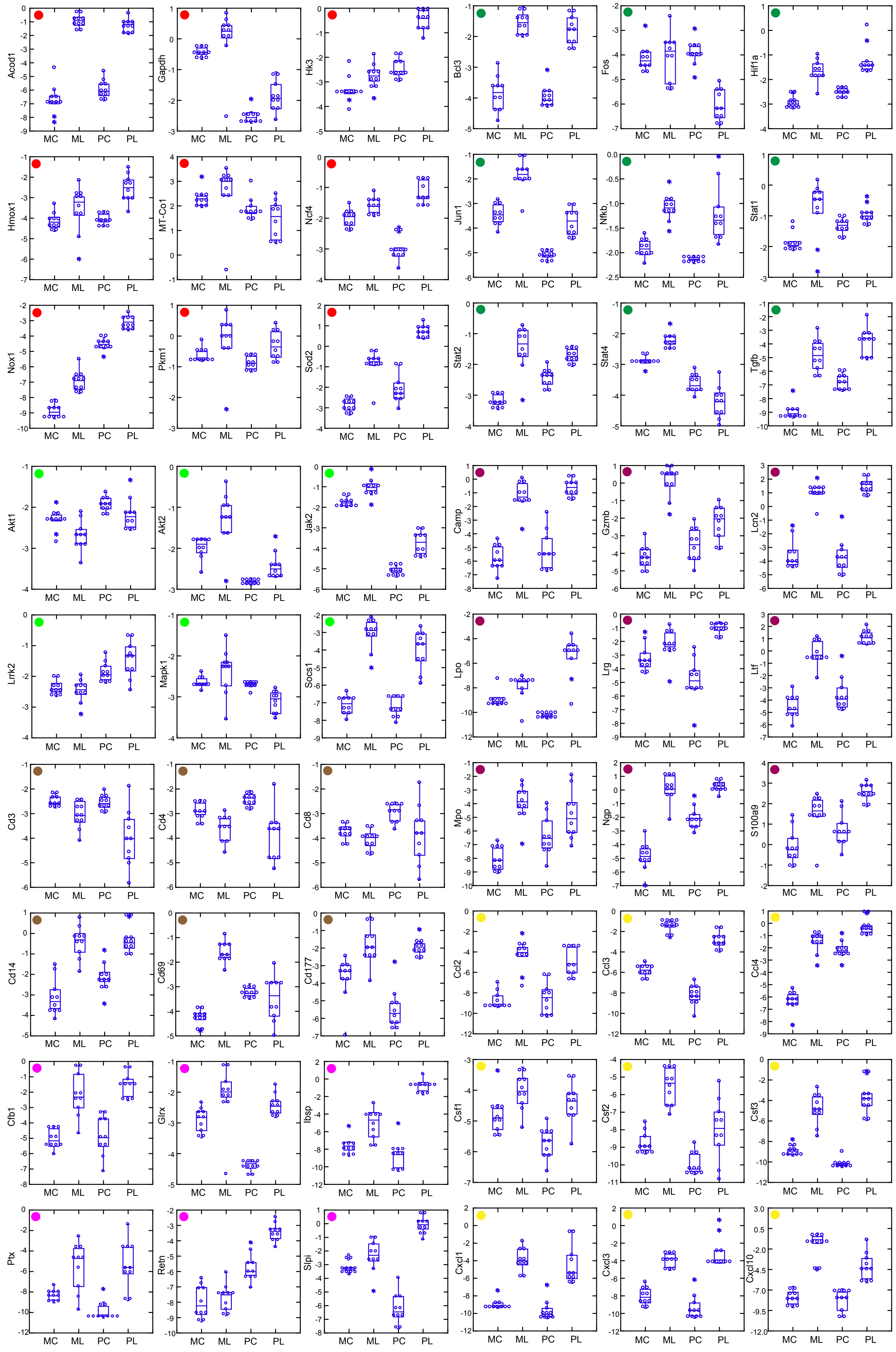

Acod1
Gapdh
Hk3
Bcl3
Fos
Hif1a
Hmox1
MT-Co1
Ncf4
Jun1
Nfkb
Stat1
Nox1
Pkm1
Sod2
Stat2
Stat4
Tgfb
Akt1
Akt2
Jak2
Camp
Gzmb
Lcn2
Lrrk2
Mapk1
Socs1
Lpo
Lrg
Ltf
Cd3
Cd4
Cd8
Mpo
Ngp
S100a9
Cd14
Cd69
Cd177
Ccl2
Ccl3
Ccl4
Cfb1
Glrx
Ibsp
Csf1
Csf2
Csf3
Ptx
Retn
Slpi
Cxcl1
Cxcl3
Cxcl10
MC
ML
PC
PL
