## Supplementary material for "The white-footed deermouse, an infection-tolerant reservoir for several zoonotic agents, tempers interferon responses to endotoxin in comparison to the mouse and rat": Figure S1

| Genus Normalization | *Mus* reads $R^2$ | *Mus* 12S rDNA $R^2$ | *Mus* Ptprc $R^2$ | *Peromyscus* reads $R^2$ | *Peromyscus* 12S $R^2$ | *Peromyscus* Ptprc $R^2$ |
|---|---|---|---|---|---|---|
| *Mus* reads | 1.000 | | | | | |
| *Mus* 12S rDNA | 0.994 | 1.000 | | | | |
| *Mus* Ptprc | 0.969 | 0.945 | 1.000 | | | |
| *Peromyscus* reads | 0.394 | 0.371 | 0.426 | 1.000 | | |
| *Peromyscus* 12S | 0.416 | 0.396 | 0.446 | 0.986 | 1.000 | |
| *Peromyscus* Ptprc | 0.424 | 0.401 | 0.457 | 0.996 | 0.990 | 1.000 |

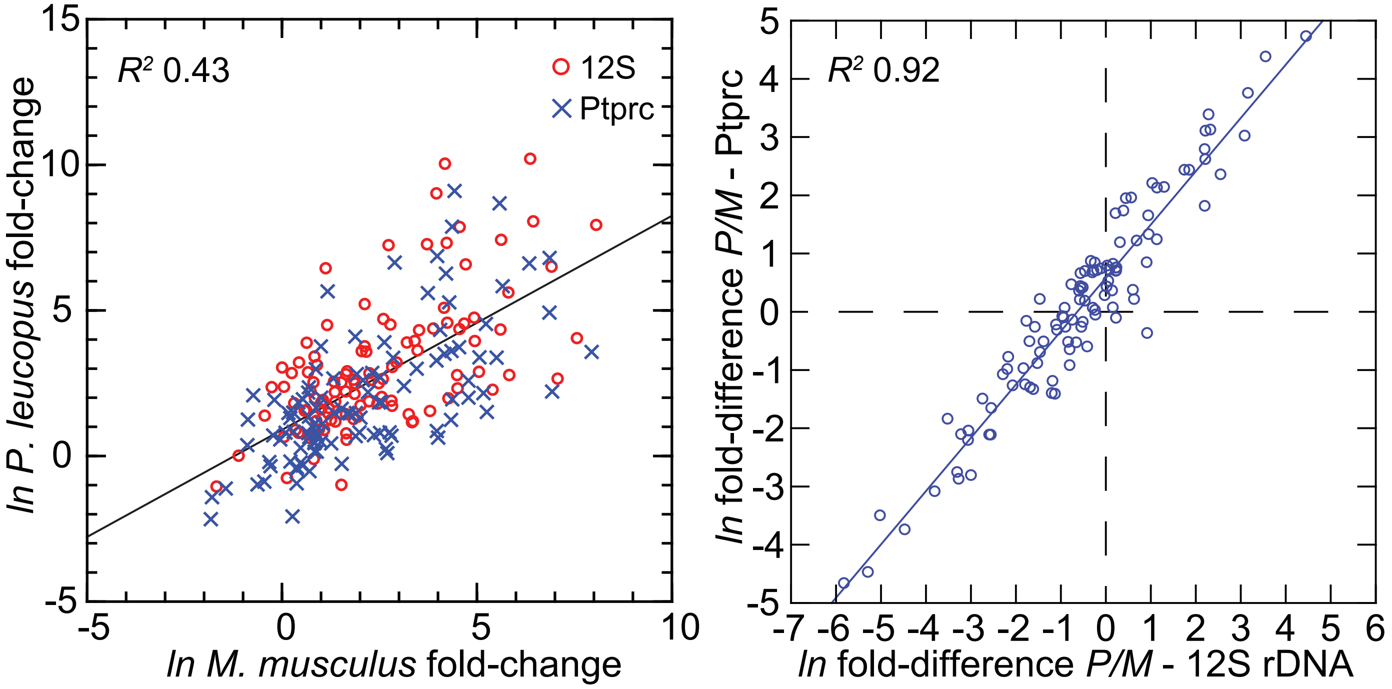


Figure S1. Comparison of three different methods for normalization for cross-species targeted RNA-seq.

The normalization options were total reads for the same sample, unique reads for the mitochondrial 12S rRNA, and unique reads for the Ptprc (CD45) transcript. The variable for comparison was, first, the mean fold-change between LPS-treated and control animals of each sex and by each of the three methods. The 109 targets and all values for the analysis are in Table S3. The coefficients of determination ($R^2$) were calculated for each of the pairs and for within each species and across species (columns B-G of Table S3). The results of this analysis are in the matrix of the upper panel. Within each species the outputs from the 3 different methods were highly similar.

Since the aim was to identify a normalization procedure suitable for use on blood sample with differing concentrations of nucleated cells and different states of activation, we next compared normalization by 12S rRNA with normalization by Ptprc. The lower left panel compares in the same scatterplot the LPS to control fold-changes by each method and the *P. leucopus* (P) result regressed on the *M. musculus* (M) for the same gene. As expected, the two species differed overall in many of their responses, and this was more pronounced for genes with the higher magnitudes of fold-change. But areas of distributions of datapoints over the space for each of the methods overlapped.

The next question was whether there would a difference in outcome for a cross-species comparison. For this we used the metric of the log-transformed ratio of the LPS:control fold- change for *P. leucopus* to the LPS:control fold-change for *M. musculus*. To avoid confusion with the “fold-change” variable, we termed this cross-species variable “fold-difference”. These values from each of two methods are in columns J and K of Table S3. If the two methods yielded commensurate results, they would be highly correlated. The lower right panel of the figure is a scatterplot with linear regression and the $R^2$ value. The slope (beta) was 0.92. The results indicated that the two normalization methods, one based on a mitochondrion gene and other on a chromosome gene, were commensurate.
